# Long non-coding RNA *LINC00152* plays an oncogenic role via targeting STAT3 and c-MYC signaling in esophageal adenocarcinoma

**DOI:** 10.1101/2020.02.19.956920

**Authors:** Honglin Zhao, Matthew Xiao, Huijie Zhao, Zhuwen Wang, Derek Nancarrow, Guoan Chen

## Abstract

**Objective:** Esophageal cancer remains a threat to public health with an increasing incidence and low survival rate worldwide. In the past thirty years the rates for esophageal adenocarcinoma (EAC) have increased over 500%. Prior studies have linked Barret’s esophagus (BE), Low grade dysplasia (LGD), and High grade dysplasia (HGD) as general precursors to EAC. However, the exact pathways by which EAC occurs have not been uncovered. Recent genomic studies have discovered a new family of active RNA species named long non-coding RNAs (lncRNAs). Of which, *LINC00152* has been linked to several human cancers and shown to promote cell proliferation in lung, gastric, hepatocellular, colorectal, and clear cell renal carcinoma. This study is to investigate the roles of *LINC00152* in EAC using EAC patient data and EAC cell lines.

**Methods:** We used *LINC00152* specific siRNAs to knockdown *LINC00152* and used the Gateway cloning method to generate stable overexpression of *LINC00152* in Flo, OE19, and OE33 cell lines for *in vitro* study. The cells were tested for changes in cell proliferation, colony formation, invasion and migration. Real-time PCR assay was used for detecting mRNA expression and Western blot was used for examining altered protein expressions affected by *LINC00152*. Data analysis were performed using excel and Prism. Statistical differences were assessed using the Student’s T-test. Survival analysis was done using Kaplan–Meier estimates.

**Results:** This study found that high levels of *LINC00152* correlated positively with tumor progression, invasive potential, and TNM stage advancement in EAC. *LINC00152* knockdown could inhibit cell proliferation, colony formation, and cell invasion. Western blot and Real-time PCR results suggests that *LINC00152* may active via STAT3 and c-MYC signaling as both demonstrated changes following knockdown and overexpression experiments.

**Conclusions:** This study indicated that *LINC00152* might be used as both a biomarker and a novel therapeutic target to improve the outcome for EAC patients. Further characterization of *LINC00152* as a novel therapeutic target for EAC is warranted.

## Introduction

Worldwide, esophageal cancer (EC) is the sixth leading cause of cancer death and is responsible for over 509,000 cases (5.3 %) [1]. In the United States, esophageal adenocarcinoma (EAC) has eclipsed squamous cell carcinoma as the predominant histologic type and usually afflicts white males. It is estimated that 13,020 American males will die from esophageal cancer in 2019. Although diagnosis has been improving, 5-year survival rate is only 19%, which is among the lowest of all cancer types [2]. These low survival rates are partly due to the fact that most EAC cases are diagnosed at a higher stage [3]. The known risk factors to EAC include gastroesophageal reflux, Barrett’s esophagus (via metaplasia-dysplasia-neoplasia sequence), obesity, Caucasian race, increasing age, alcohol and smoking. Because multiple molecular pathways converge to trigger unregulated growth, invasion, and metastasis in EAC, the exact mechanisms are not fully understood. Thus, there is an urgent need for new bio-molecular markers that may aid in the early diagnosis or therapy [4].

The human genome contains only 20,000 protein-coding genes, representing less 2% of the total genome, whereas a substantial fraction of the human genome can be transcribed, yielding many short or long intervening/intergenic noncoding RNAs (lncRNAs) with limited protein-coding capacity [5]. Although most human lncRNAs have been identified, less than 1% of those have been characterized [6,7,8,9]. Recent studies on the functionality of lncRNAs in the meshwork of cancer pathways, have found lncRNAs to have both oncogenic and tumor suppressing functions in regulating tumorigenesis [10].

Of the studied lncRNAs, *LINC00152* has been linked to several human cancers and promoted cell proliferation, invasion, metastasis, and apoptosis in lung, gastric, hepatocellular, colorectal, gallbladder, and clear cell renal carcinoma [11,12,13,14]. High *LINC00152* expression levels were associated with chemo-resistance as well as poor prognosis. Additionally, studies also suggested that *LINC00152* might be involve in the carcinogenesis of several cancers by disturbing various signaling pathways [15]. Thus, *LINC00152* may be used as a new tumor molecular biomarker, applicable for tumor diagnosis, targeted therapy, and prognosis assessment [16]. However, the expression and biological roles of *LINC00152* in EAC remains unclear. This study tried to determine the role of *LINC00152* in EAC and identify the molecular functions, regulatory mechanisms, and clinical applications.

## Results

### Overexpression of *LINC00152* is correlated with tumor progression in patient with EAC

To investigate the *LINC00152* expression status and its diagnostic potential in esophagus adenocarcinoma, we performed expression analysis with RNA-Seq data from the UM cohort[17] (University of Michigan BEs 25, LGDs 8, HGDs 25 and EACs 11). We have found *LINC00152* was highly expressed in EACs as compared to precursors, and *LINC00152* was gradually increased from pre-cancer progression tissues to EACs (Fig. 1A). We next evaluated the association of *LINC00152* and patient survival. Kaplan-Meier survival curves and log-rank tests showed that higher expression levels of *LINC00152* has a poor patient survival trend (p = 0.17) in UM data[17] (Fig. 1B). Non-parametric Kruskel-Wallis testing indicated that higher *LINC00152* expression was presented in poorly differentiated, more lymphocytic invasived and highly desmoplastic responded tumors (Fig. 1C-E). All these results indicated that *LINC00152* may be potentially used as a novel diagnostic marker for EAC.

**Figure 1.**
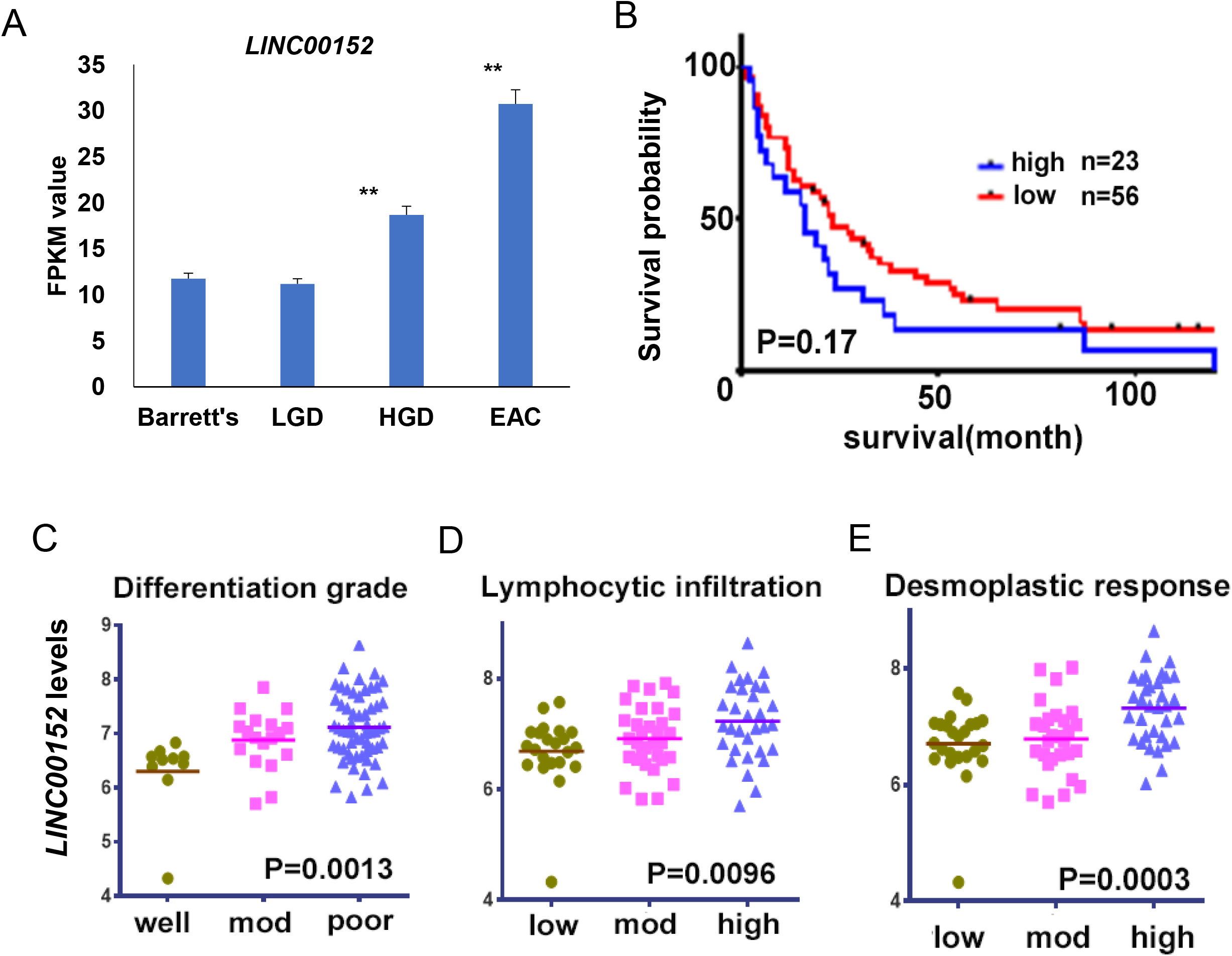
**A.** Expression of *LINC00152* increases in EAC compared to the pre-cancer progression tissues: Barrett’s Esophagus, LGD, and HGD. **B.** Kaplan-Meier curve indicated higher LINC00152 expression was unfavorable for patient survival. **C**, **D** and **E**. Higher LINC00152 expression showed poor differentiation grade, stronger lymphocytic invasion, high desmoplastic response.

### *LINC00152* knockdown decreased cell proliferation and colony formation in EAC cells

In order to perform the functional study, we first examined the expression of *LINC00152* on the 3 EAC cells (OE19, FLO, and OE33) by RT-PCR. We found that the FLO cell was the highest expression for *LINC00152*, and OE19 was the lowest (data not shown). To minimize the possibility of siRNA off-target effects, we used SMART pool gene specific siRNAs whose knockdown efficiency was greater than 80–90% as determined by qRT-PCR analysis (Fig. 2A). To explore the oncogenic role of *LINC00152*, we examined the cell proliferation status measured by WST-1 after *LINC00152* knockdown by siRNA in 3 EAC cell lines. A significant decrease in cell proliferation rate (OE19 16%, FLO 13% and OE33 38%) was found in EAC cells (Fig. 2B). *LINC00152* significantly affected cell invasion using Boyden chamber matrix assays in FLO (Fig. 2C and D). In colony formation assay, the number of colonies was markedly reduced after *LINC00152* siRNA knockdown in OE19, FLO and OE33 cells (Fig. 2E and F).

**Figure 2.**
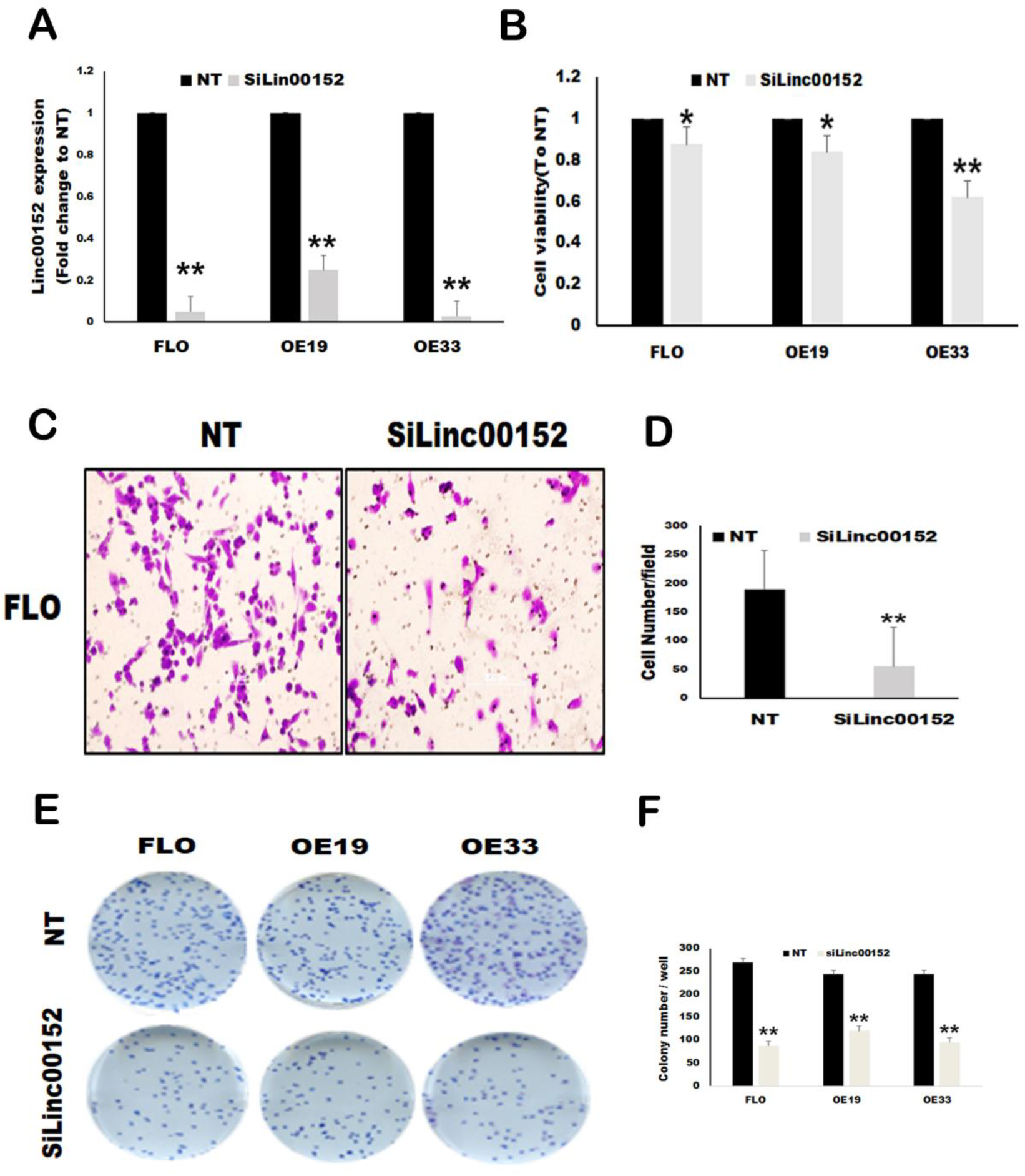
**A.** siLINC00152 RNA decreased LINC00152 expression by more than 90% in Flo-1 OE19 and OE33 cell lines, as measured by RT-PCR. **B.** Cell proliferation of EAC cell lines was measured by WST-1 assay to determine the effect on cell viability after siLINC00152 transfection for 96 hours. **C** and **D**. Invasion was decreased after *LINC00152* siRNA transfection on Flo-1. **E** and **F.** Colony formation was reduced when siLINC00152 was transfected into Flo-1, OE19 and OE33 cells.

### STAT3 and c-MYC were regulated by *LINC00152* in EAC cells

To provide molecular mechanistic insight of *LINC00152* in regulating EAC cell proliferation, we performed Western blot analysis using different antibodies covering the most of the cancer-related pathways such as STAT3, c-MYC, c-MET, p21, p27 etc. after *LINC00152* siRNA treatment in OE19, FLO and OE33 cell lines. The levels of total STAT3, phosphonate STAT3 and c-MYC proteins were found to be decreased when *LINC00152* was knocked down by siRNA at 72 hours (Fig. 3A and B), especially in OE33 and FLO, which have the higher expression of STAT3. We also performed the mRNA expression of these 2 genes and found that the mRNAs of STAT3 and c-MYC were decreased by 30–40% in both cells (Fig. 3C and D). These results indicated that *LINC00152* regulates STAT3 and c-MYC possibly through transcriptional regulation. The mechanism of these genes regulated by *LINC00152* is warranted to be analyzed in the future.

**Figure 3.**
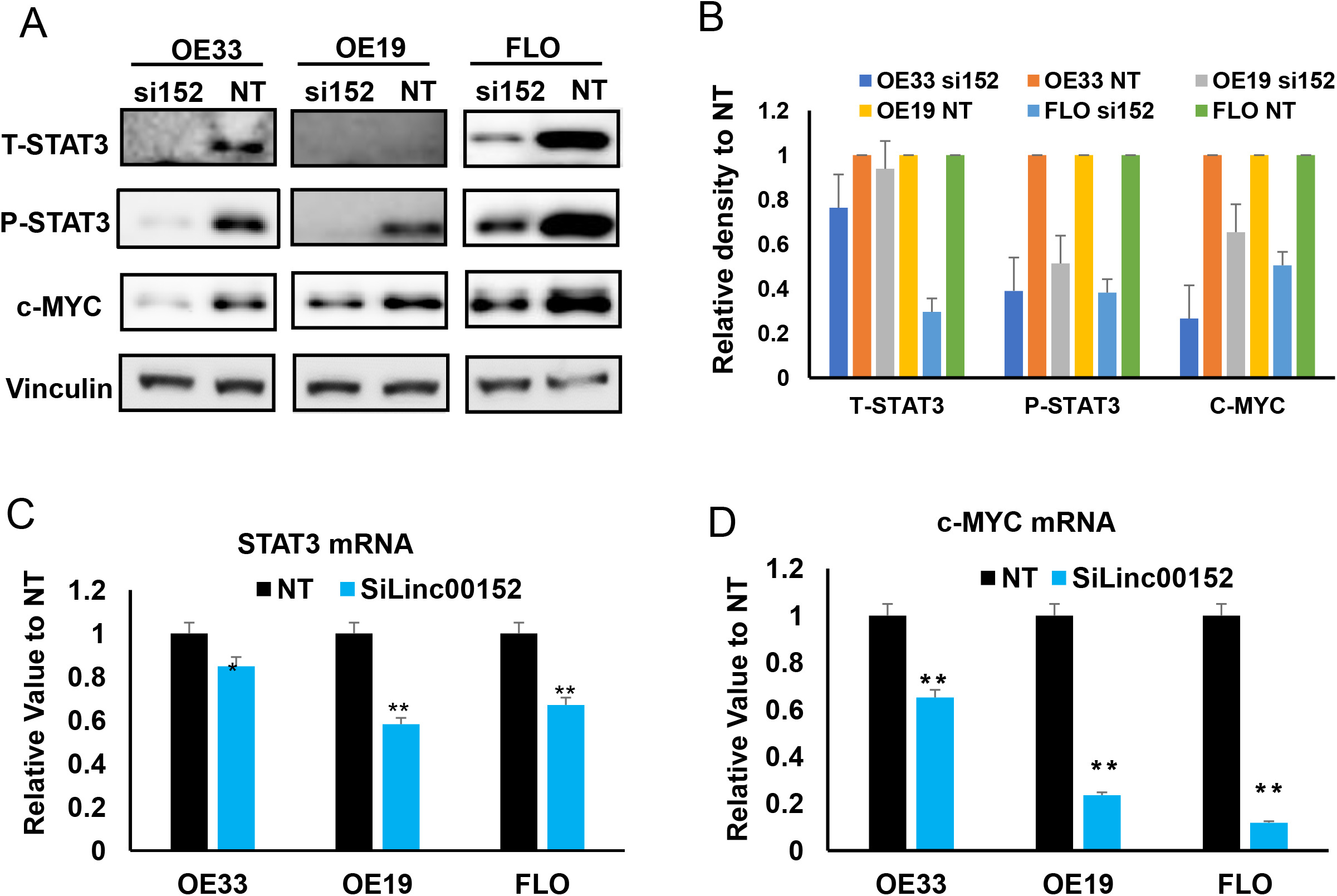
**A**. Western blot showing the altered protein expression of STAT3, c-MYC after *LINC00152* knockdown with siRNA. **B.** Quantitative analysis of image in A using ImageJ sofware. Vinculin was used as a protein loading control. Each individual band was divided by Vinculin first, then divided by NT. So each protein value (siRNA treated) was relative to their NT (NT = 1). **C** and **D.** Quantitative real time RT-PCR confirmed STAT3, c-MYC change in mRNA level.

### Overexpressed *LINC00152* confirmed in STAT3 and c-MYC pathways

To further confirm the results of *LINC00152* in EAC, we performed overexpression assay using Gateway cloning method to generate stable overexpression of *LINC00152* in OE33. Cell proliferation was significantly increased in OE33 cells after *LINC00152* overexpression (Fig. 4A). RT-PCR results confirmed the STAT3 and c-MYC upregulated accompanying *LINC00152* overexpression in mRNA level (Fig. 4B). Overexpression of *LINC00152* significantly promoted cell invasion using Boyden chamber matrix assays in OE33 (Fig. 4C and D).

**Figure 4.**
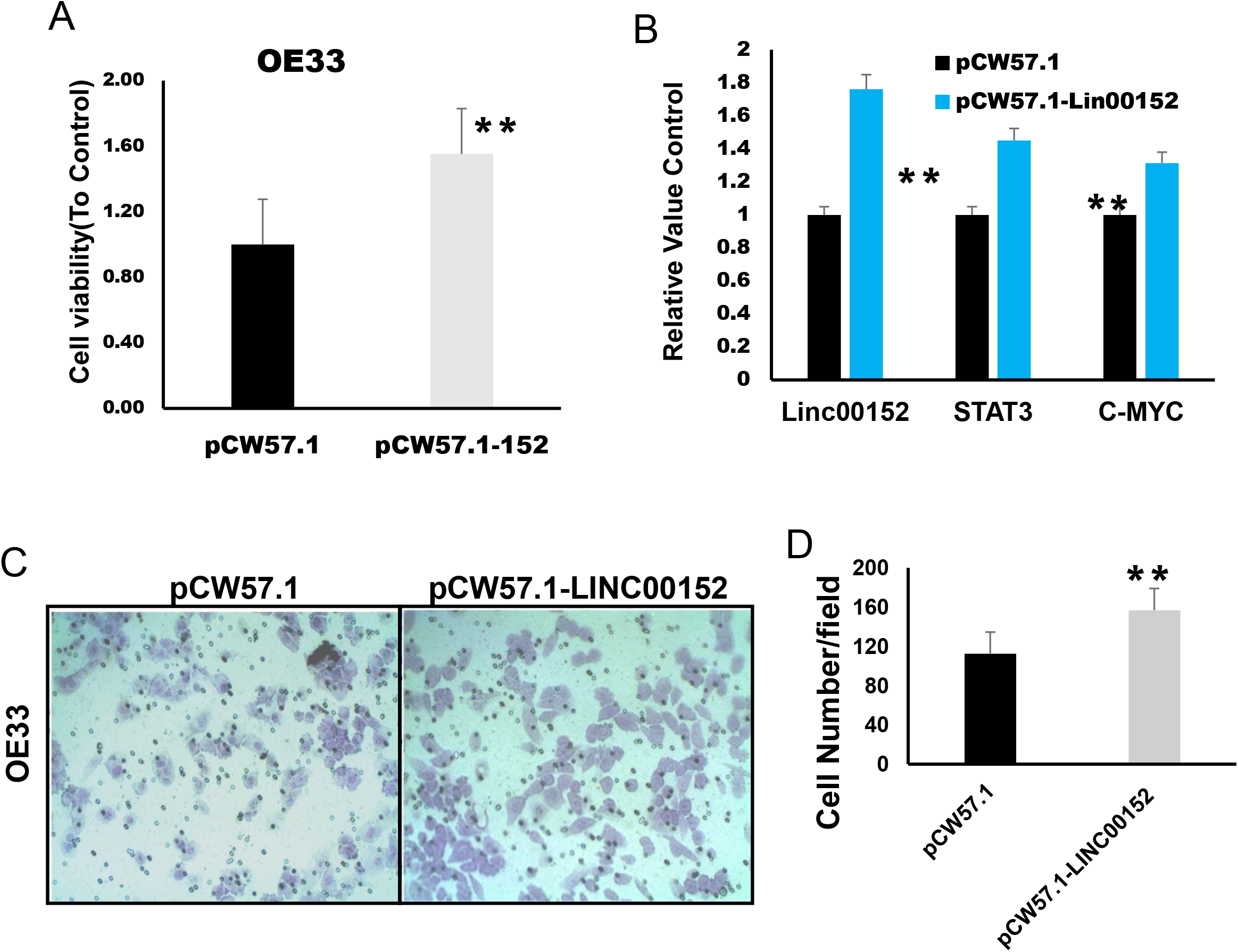
**A.** OE33 cell transfected with pCW57.1-*LINC00152* showed higher LINC00152 expression (as expected) as well as increased cell proliferation. **B.** Quantitative real time RT-PCR confirmed LINC00152, STAT3 and c-MYC change in RNA level after OE33 cell transfected with pCW57.1-*LINC00152* **. C** and **D.** OE33 invasion increased relative to the pCW57.1 vector alone

## Discussion

As highly tissue-specific drivers of cancer phenotypes, lncRNAs are potentially prime targets for cancer therapy. Many studies have confirmed their utility as biomarkers not only in cancer diagnosis but also for patient prognosis across a variety of cancers. *LINC00152* has been reported to be highly expressed in ESSC [18]. In the present study, we found that the average levels of *LINC00152* in EAC tissues were significantly higher than precursors, and higher expression of *LINC00152* was associated with a poor tumor differentiation and stronger lymphocytic invasion. These results were consistent with previous studies [19], which suggest that *LINC00152* may be potentially useful as a biomarker for EAC diagnosis and an indicator of poor survival. We found that knockdown of *LINC00152* suppressed tumor cell proliferation and colony formation capability of EAC cells, and also affected tumor cell invasion, which was consistent with the results observed in lung cancer [11].

LncRNAs may regulate oncogenesis by different molecular mechanisms in different cancers. Previous study observed that *LINC00152* can specifically recognize the EGFR-binding site and activate the PI3K/AKT signaling pathway to promote the proliferation of gastric cancer cells [20]. In this study, *LINC00152* knockdown leads to reduced cell growth-related proteins STAT3 and c-MYC in EAC, whereas increased STAT3 and c-MYC expression after *LINC00152* upregulation. This suggests that the molecular signaling affected by *LINC00152* in EAC may be different from other cancers. STAT3 was shown directly or indirectly upregulating the expression of genes required for uncontrolled proliferation and survival, including the genes that encode c-MYC. The increased expression of STAT3 may also positively affect c-MYC, thus precipitating a series of events during oncogenesis including cell proliferation, angiogenesis, and anti-immune responses [21]. We found that STAT3 mRNA and protein were decreased after *LINC00152* knockdown indicating that *LINC00152* regulated STAT3 possibly at the transcription level. Thus, *LINC00152* mainly acted through STAT3 and c-MYC pathways to influent EAC progression (Figure 5).

**Figure 5.**
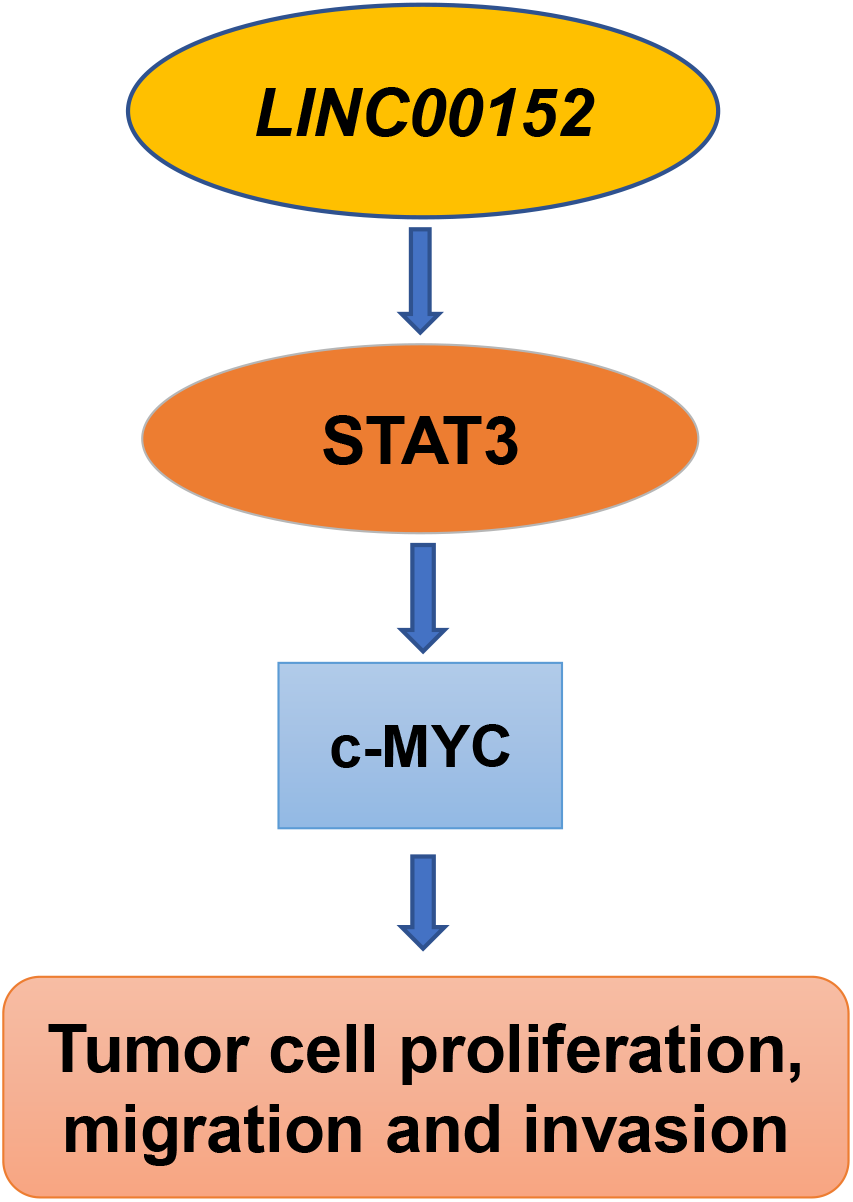
Schematic model of the role of *LINC00152* in EAC. LINC00152 affected EAC cells proliferation and invasion may via STAT3 and c-MYC signaling.

In conclusion, we found that higher expression of *LINC00152* was closely associated with high malignancy of EAC. We also demonstrated the functions of *LINC00152* in promoting proliferation, invasion, and migration of EAC in vitro. The underlying mechanisms may involve STAT3 and C-MYC pathways.

## Materials and Methods

### Patient and Tissues

All samples were obtained after receiving written, informed patient consent according to the approval and guidelines of the University of Michigan institutional review board. Sample tissues were taken from esophagectomy patients who had not received preoperative radiation or chemotherapy. All of the samples were taken between 1992 and 2015. Each specimen was immediately frozen in liquid nitrogen and stored at −80°C until further use.

### Cell Lines and Cell Culture

The Flo cell line used in this experiment was derived from a stage IIb EAC patient in the laboratory and has been previously described [3]. OE19 and OE33 were derived from stage IIa and III EAC patient tumors, respectively, and are both maintained by The European Collection of Cell Cultures (Sigma-Aldrich). Cell lines were either grown in DMEM (Flo) or RPMI-1640 (OE19 and OE33) supplemented with 10% fetal bovine serum (FBS; Atlanta Biologicals) and 1% penicillin/streptomycin/fungizone (Invitrogen). Cells were allowed to grow in an incubator at 37°C in 5%CO_2_/95% air. Prior to treatment, cells were plated on 6-well plates at a density of 150,000 cells/well in 2 mL of medium.

### RNA Isolation and Real-time PCR (qRT-PCR)

Total RNAs were isolated from the treated cell lines using the QIAGEN RNeasy Mini Kit (Qiagen) and treated with DNase I according to the manufacturer’s instructions. The concentration for the RNAs was quantified using the Nanodrop 3000 (Nanodrop Products). Total 2ug RNAs were reverse transcribed using High Capacity cDNA Reverse Transcription Kit (Thermo Fisher Scientfic). Real-time PCR was performed on the cDNA using the Platinum SYBR Green PCR Master Mix (Invitrogen) on the ABI Prism 7900HT Detection System (Life Technologies) through the University Michigan Sequencing Core. The PCR amplification was performed for 40 cycles of 95°C for 15 second and 60°C for 60 second and Significant differences of relative quantification were determined using the 2-ΔΔCt method [22] with normalization to GAPDH or β-actin. The forward primer sequence for LINC00152 was: 3’-TCTTCACAGCACAGTTCCTGG-5’. and the reverse primer sequence was 3’-GGCTGAGTCGTGATTTTCGG-5’. The forward primer sequence for GAPDH was: 3’-CTCTGCTCCTCCTGTTCGAA-5’ and the reverse sequence was 3’-ACGACCAAATCCGTTTGACT-5’. The forward primer sequence for β-actin was: 3’-CTCTTCCAGCCTTCCTTCCT-5’ and the reverse sequence was 3’-AGCACTGTGTTGGCGTACAG −5’. The forward primer sequence for STAT3 was: 3’-TGGCCCAATGGAATCAGCTAC-5’ and the reverse sequence was 3’-CTGCTGGTCAATCTCTCCCA-5’. The forward primer sequence for c-MYC was: 3’-CAGCGACTCTGAGGAGGAAC-5’ and the reverse sequence was 3’-TGTGAGGAGGTTTGCTGTGG-5’.

### Protein Isolation and Western Blot

Total cellular protein was extracted using 50-80 μL of a prepared lysis buffer (150 mM NaCl; 20 mM Tris, pH 7.5; 1 mM EDTA; 1 mM EGTA; 2.5 mM Na4 P2 O7; 1 mM glycerol 2-phosphate disodium salt hydrate; 1 mM Na3 VO4; 1% Triton X-100) supplemented with Protease Inhibitor Cocktail (20 μL/1 mL lysis buffer; Cat#P8340 Sigma-Aldrich). Lysis buffer was allowed to react for 30 minutes on ice before the mixture was centrifuged at 14,000 rpm and 4°C for 20 minutes. The protein concentration was then quantified using the DC Protein Assay Kit II (Bio-Rad), and the absorbance was read at 750 nm using the FLx800 Fluorescence Microplate Reader (BioTek). To denature the protein, 10 to 20 μg of protein were boiled in 1x LDS Non-Reducing Sample Buffer (Life Technologies) and supplemented with 5% 2-mercaptoethanol (Bio-Rad Laboratories) for 8 minutes at 100 °C. The samples were then run on 10% Tris-glycine gels (Invitrogen), and transferred to 0.45um PVDF membrane (Millipore) for 2.5 hours on ice. The membranes were blocked with Blotting-grade Blocker (Bio-Rad) for 1 hour at room temperature, and then hybridized overnight with primary antibody at 4°C and washed three times using TBST buffer. The primary antibody STAT3, c-Myc, and venculin were purchase from Cell Signaling. The primary antibodies were made at 1:1,000 dilution in 5% milk TBST buffer. Next, the membrane was incubated with secondary antibody for 1 hour at room temperature and washed three times, and then detected using the Amersham ECL Prime Western Blotting Detection Reagent (GE Healthcare Life Sciences) by ChemiDoc MP imaging System (Bio-Rad).

### Small Interfering RNA (siRNA) Transfection

siRNAs targeting LINC00152 were purchased from Dharmacon and were pooled. The siRNA was transfected at a concentration of 10nM for cell migration, invasion, Western blotting and qRT-PCR experiments. Lipofectamine RNAiMAX Reagent (Thermo Fisher Scientific) was used according to the manufacturer’s instructions. A non-targeting siRNA#1 (Dharmacon) was used as a control and was transfected at 10nM for all experiments. In addition, a clonogenic assay was performed. A six well plate was used with 250 cells per well. The colonies were allowed to grow for two weeks before they were stained with crystal violet and counted. Data was collected in triplicate.

### Cell Proliferation Assays

Cells were first plated on a 96-well plate at a concentration of 1000 cells/well. They were treated with either siRNA or control siRNA #1. After 24 hours, the medium was changed, Cell proliferation was assessed using the WST-1 (Roche) assay according to the manufacturer’s instructions. Cell proliferation was quantified using the ELx808 Absorbance Microplate Reader (BioTek Instruments, Inc.) at 450 nm with a background wavelength at 630 nm. The readings were collected 72 hours after treatment. Data was collected in triplicate.

### Matrigel Invasion and Migration Assay

For the invasion experiments, ice-cold Matrigel (BD Bioscience) was thawed on ice and combined with coating buffer until a concentration of 80ug/mL was reached. Afterwards, 100 uL of the Matrigel and coating buffer mix was added to the upper chamber of a 24 well invasion chamber system (BD Bioscience). 1×10^6 cells were plated in the upper chamber. These cells were placed in DMEM medium without FBS while 20% FBS was added to the medium of the lower chamber to serve as a chemoattractant. After 48 hours, non-invasive cells and the Matrigel on the bottom of the chamber were removed using a cotton swab, and the cells beneath the chamber were stained with crystal violet. Assays were performed in triplicate, and cells were counted at a magnification of 4x. Migration was measured using the same procedure without Matrigel.

### Construct of *LINC00152* Overexpression Stable Cell Line

A *LINC00152* mammalian expression construct was purchased from Dharmacon and was inserted into lentivirus vector pCW57.1 in UM Vector Core Facility by using Gateway recombination cloning system and stably transfected into OE33 cell line.

### Affymetrix Microarray and RNAseq Analysis

Affymetrix Microarray and RNAseq analysis were performed by The University of Michigan DNA Microarray Core Facility[17].

### Statistical Analysis

The Kaplan-Meier survival test was calculated using the GraphPad Prism7 software and p-values were determined by log-rank test. Means, standard deviations, and figures were all calculated and created using Microsoft Excel software. The error bars presented in the figures represent the standard deviation of the samples. Student’s T-tests and correlation coefficients were applied for all necessary experiments and p-value of <0.05 was used.

## Acknowledgments

We thank Daysha Ferrer-Torres, Jules Lin, Andrew Chang and David G. Beer for helpful information and comments.

